# Evaluating the Roles of Drift and Selection in Trait Loss along an Elevational Gradient

**DOI:** 10.1101/2024.06.12.598645

**Authors:** Sophia F. Buysse, Samuel G. Pérez, Joshua R. Puzey, Ava Garrison, Gideon S. Bradburd, Christopher G. Oakley, Stephen J. Tonsor, F. Xavier Picó, Emily B. Josephs, Jeffrey K Conner

## Abstract

Traits that have lost function sometimes persist through evolutionary time. These traits may persist if there is not enough standing genetic variation for the trait to allow a response to selection, if selection against the trait is weak relative to drift, or if the trait has a residual function. To determine the evolutionary processes shaping whether nonfunctional traits are retained or lost, we investigated short stamens in 16 populations of *Arabidopsis thaliana* along an elevational cline in northeast Spain. We found a cline in short stamen number from retention of short stamens in high elevation populations to incomplete loss in low elevation populations. We did not find evidence that limited genetic variation constrains the loss of short stamens at high elevations, nor evidence for divergent selection on short stamens between high and low elevations. Finally, we identified loci associated with short stamens in northeast Spain that are different from loci associated with variation in short stamen number across latitudes from a previous study. Overall, we did not identify the evolutionary mechanisms contributing to an elevational cline in short stamen number but did identify different genetic loci underlying variation in short stamen along similar phenotypic clines.

**Teaser text:** The evolutionary mechanisms underlying loss or retention of traits that have lost function are poorly understood. Short stamens in *Arabidopsis thaliana* provide a compelling system to investigate the roles of genetic drift and selection in trait loss across latitudinal and elevational clines. This study investigates how drift and selection shape short stamen loss in 16 populations of *A. thaliana* along an elevational gradient in Northeast Spain. An investigation of the loci underlying variation in short stamen number suggests variants in different genes may cause trait loss in similar phenotypic clines within a species.

## Introduction

Traits that have lost function are often lost through evolutionary time, yet some persist. Nonfunctional traits could be lost through direct selection against the trait (Dorken et al., 2004; Lahti et al., 2009), correlated responses to selection on other traits caused by pleiotropy or linkage disequilibrium (Yoshizawa et al., 2012), or the accumulation of selectively neutral mutations (Fong et al., 1995). In contrast, nonfunctional traits may be retained by evolutionary constraint if the nonfunctional trait is genetically correlated with a different, functional trait (Lande, 1979; Walsh & Blows, 2009). Nonfunctional trait loss could also be prevented if there is not enough standing genetic variation for selection to act on the trait (Lahti et al., 2009) or if weak selection against the trait is unable to overcome drift (Charlesworth, 2009). Both processes may be pronounced in species with small effective population sizes. Determining the evolutionary processes shaping whether nonfunctional traits are retained or lost will have broad implications for our understanding of the interplay of direct selection, correlated responses to selection, and genetic drift (reviewed in Futuyma, 2010).

Here, we use an elevational cline in short stamen number in *A. thaliana* from Northeast Spain to investigate how evolutionary processes interact to shape variation in a trait that has lost function. Flowers across almost all of the 3,700 species in the Brassicaceae family have four long and two short stamens (Zomlefer, 1994). This tetradynamous stamen arrangement is strongly phylogenetically conserved. The function of short stamens in Brassicaceae is unknown, especially in self-pollinating species. In outcrossing species, the presence of short stamens increased the duration of pollinator visits (Kudo, 2003) and short stamen anthers produce more pollen though less of it is removed than pollen from long stamen anthers during pollinator visits (Conner et al., 1995). There is also selection to maintain both long and short stamens in *R. raphanistrum*, an outcrossing species (Conner et al., 2003; Waterman et al., 2023), but the function of short stamens even in outcrossing species is unknown. Loss of short stamens, resulting in flowers with four long stamens and either one or zero short stamens (Fig. 1A), has been observed in self-pollinating *Arabidopsis thaliana* and a few other Brassicaceae genera (Bowman et al., 1999; Matsuhashi et al., 2012; Müller, 1961) but is not common across the family and has not been observed in *Arabidopsis lyrata* or *Capsella bursa-pastoris*, close relatives of *A. thaliana*. See Royer et al. (2016) for a review of stamen number variation.

**Figure 1:**
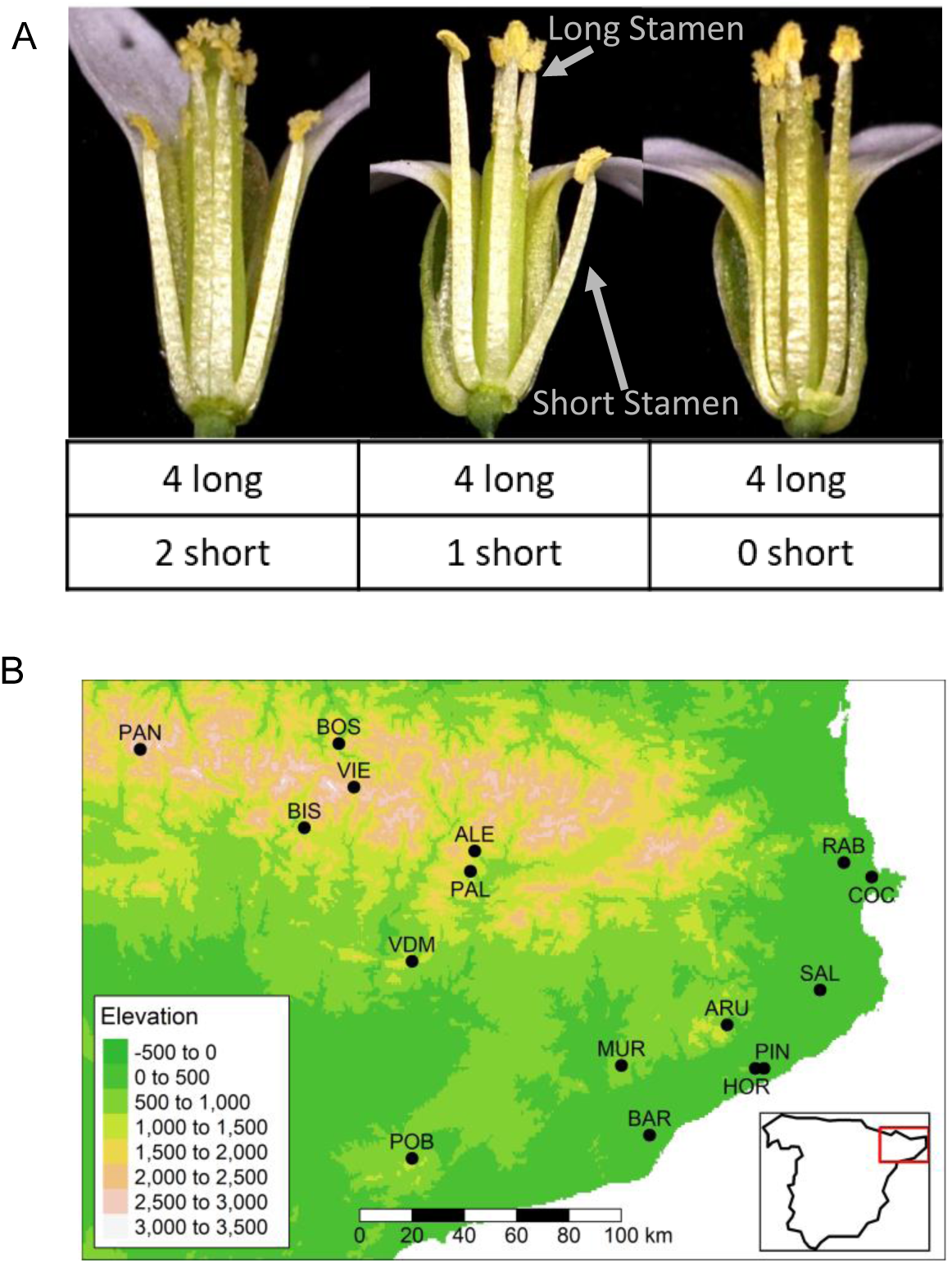
Populations studied across an elevation gradient. **A)** Natural variation in short stamen number. Flowers have 2 (left), 1 (center), or 1(right) short stamens. All flowers have four long stamens. Photos by Frances Whalen. **B)** Each point represents the seed collection site for 16 populations of *A. thaliana* across northeast Spain. The map coloring indicates elevation from green at low elevation to white at high elevation. **Alt text:** Photos of three Arabidopsis flowers and a map showing seed collections sites in Northeastern Spain.

*A. thaliana* evolved to be almost entirely self-fertilizing from an outcrossing ancestor between 0.5 – 1 million years ago (Durvasula et al., 2017; Tang et al., 2007). In flowering plants, a transition from outcrossing to self-pollination and subsequent relaxation of selection for pollination often results in a suite of trait changes, called the ‘selfing syndrome’, that includes smaller flowers and reduced distance between the anthers and stigma (Sicard & Lenhard, 2011). Self-pollination also decreases local effective population size (Caballero, 1994), increasing the effects of genetic drift and thus reducing within-population genetic variation.

Because short stamens in *A. thaliana* are below the stigma, we expect them to have little or no function in self-pollination. Prior work confirmed these expectations: short stamens in *A. thaliana* do not contribute significantly to self-fertilized seed number and many populations in the native range have flowers with zero or one short stamens (Royer et al., 2016). Almost all flowers have four long stamens, and the short stamens are not reduced but rather lost entirely. Fixation for zero short stamens has not been observed in all *A. thaliana* natural populations studied to date (Royer et al., 2016). However, variation between populations is common. For example, southern European populations are more likely to lose short stamens while northern European populations typically retain both short stamens. Royer et al. (2016) identified three QTL underlying the variation in short stamen number across Europe.

Short stamen loss in *A. thaliana* is an excellent model to study the evolutionary mechanisms influencing variation in a trait that has lost most or all function. Because northern European populations of *A. thaliana* have undergone repeated bottlenecks that reduced effective population size, and thus standing genetic variation (Beck et al., 2008; François et al., 2008; Lewandowska-Sabat et al., 2010), short stamens may be retained in these populations because there is insufficient genetic variation for stamen number and/or weak selection against short stamens cannot overcome drift. Alternatively, the latitudinal cline of the European populations could result from local adaptation for short stamen number or another correlated trait.

There are many evolutionary scenarios that could result in phenotypic clines, but the same evolutionary processes occurring across latitudes may shape elevational clines in short stamen number. High elevations were glaciated, even at southern latitudes (Hughes et al., 2006) and while Northeast Spain does include putative refugia regions (Médail & Diadema, 2009), prior work in Northeast Spain has shown high elevation montane populations have less genetic variation and smaller effective population sizes than lower elevation coastal populations (Gomaa et al., 2011; Montesinos et al., 2009). Further, because environmental factors vary similarly with latitude and elevation, selection may result in similar phenotypic clines. Yet, latitudinal and elevational phenotypic clines are not always parallel (Daco et al., 2021; Kooyers et al., 2015). Even if the same evolutionary processes are resulting in parallel phenotypic clines, they may not have a parallel genetic basis (Fulgione et al., 2022; Gamba et al., 2024). If drift contributes to the phenotypic clines, a parallel genetic basis is unlikely as drift acts equally and randomly across the genome and any gene in the short stamen developmental pathway could be impacted. If selection contributes to the phenotypic clines, a parallel genetic basis is still unlikely to result from standing genetic variation for short stamens in *A. thaliana* due to small effective population sizes, no evidence for strong selection on short stamens, and the multiple loci involved in the latitudinal cline (MacPherson & Nuismer, 2017). Thus, we do not expect a parallel genetic basis for short stamen loss across latitude and across elevation despite a strong expectation for parallel phenotypic clines.

To understand the processes shaping variation in short stamen loss, we assessed 16 populations of *A. thaliana* along an elevational gradient in Northeast Spain. We counted short stamen number on plants grown in a growth-chamber common garden and sequenced multiple individuals per population to quantify genetic variation. We identified an elevational cline in short stamen number in Northeast Spain, with more stamen loss at low elevation, similar to the previously described latitudinal cline (Royer et al., 2016). We then used multiple regression, polygenic adaptation detection, and genome wide association studies to address possible evolutionary mechanisms contributing to the cline:

**1)** Does variation in effective population size contribute to the short stamen loss cline? Less stamen loss in populations with less genetic variation after accounting for variation due to elevation would be consistent with the hypothesis that stamen loss is constrained by a lack of genetic variation and less effective selection.
**2)** Does local adaptation contribute to the short stamen loss cline? Evidence for divergent selection on short stamen number between high and low elevation populations would support local adaptation contributing to the cline.
**3)** Are the same loci associated with variation in short stamen number across elevation as across latitude? Identifying loci within the three previously identified QTL (Royer et al., 2016) would support parallel genomic evolution between the latitudinal and elevational clines in short stamen number. Alternatively, we may detect different large-effect loci or small effect loci, many of which will be below our detection threshold.

## Materials and Methods

### Seed Collection

Population collection sites span a continuous elevation gradient from 61m to 1706m above sea level along the Mediterranean coast and into the Pyrenees (Fig. 1B; Fig. 2-3 legend; Table S1). Seeds from five to nine individuals were haphazardly collected from multiple patches in each of 16 populations (*N* = 112 genotypes) in northeast Spain (Montesinos-Navarro et al., 2011). These lineages are also used in Montesinos-Navarro et al. (2011), Montesinos-Navarro et al. (2012), and Wolfe and Tonsor (2014). They include the 10 populations from Montesinos et al. (2009) and the 9 populations from Gomaa et al. (2011).

**Figure 2:**
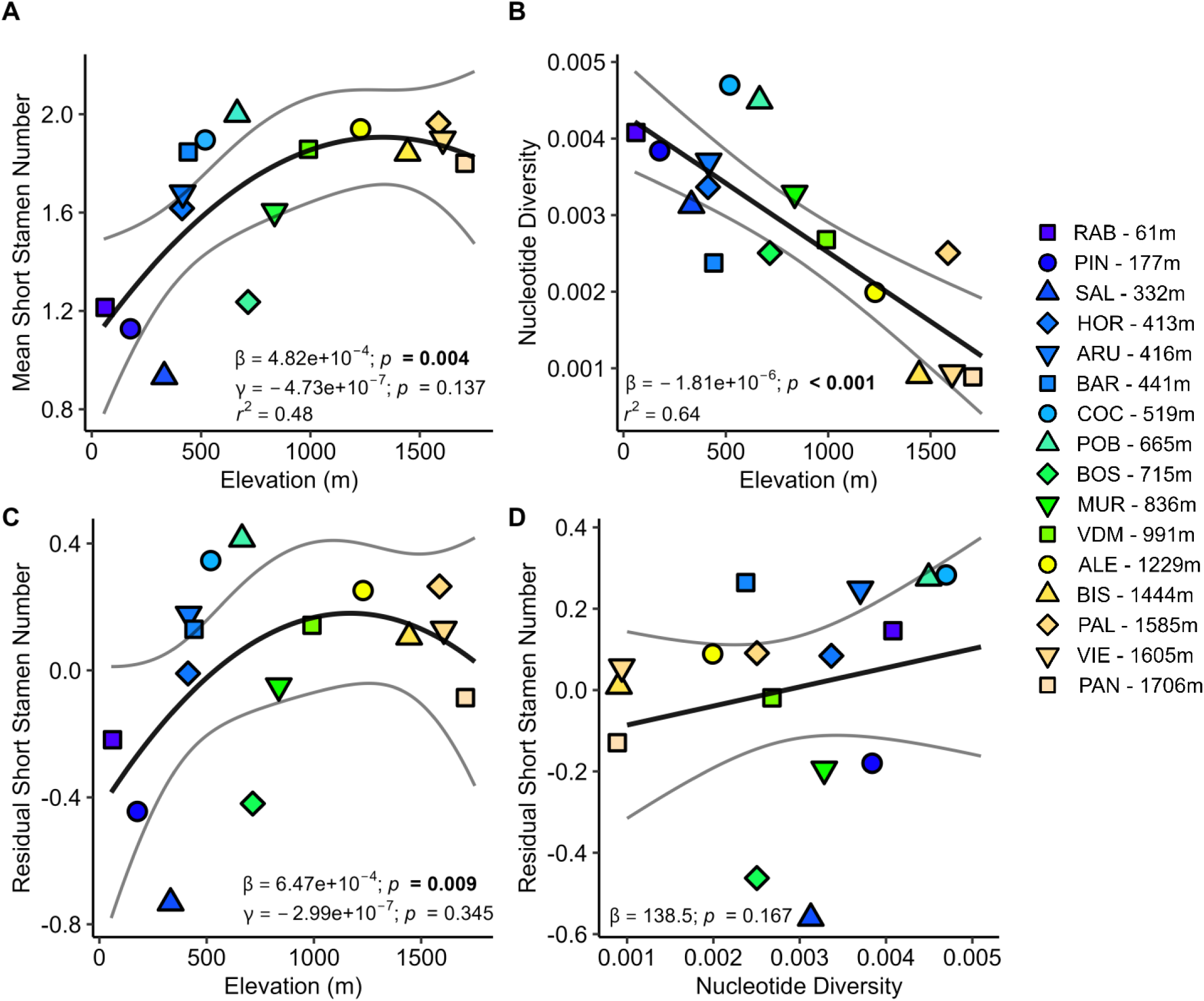
Effective population size does not explain retention of short stamens at high elevation. **A)** Mean short stamen number shows a quadratic elevational cline. **B)** Elevation strongly predicts population mean pairwise nucleotide diversity, our measure of effective population size. **C)** The residuals of the model of short stamen number regressed on nucleotide diversity then regressed on elevation and **D)** the residuals of the model of short stamen number regressed on quadratic elevation then regressed on nucleotide diversity. C and D visually underestimate the confidence intervals because they do not include uncertainty in calculating the residuals; the interpretation remains the same because statistics displayed on the figures are from the full model (*r*^2^ = 0.50; Table S3). In all panels, the color of each point represents the population elevation, black lines are the regression, and grey lines are 95% confidence intervals. **Alt text:** Graphs depicting relationships between short stamen number, elevation, and nucleotide diversity. Short stamen number increases with elevation and the linear term is significant but the quadratic term is not. Nucleotide diversity decreases with elevation. After accounting for nucleotide diversity, short stamen number still increases with elevation. After accounting for elevation, nucleotide diversity now increases with elevation though the slope is not significant.

**Figure 3:**
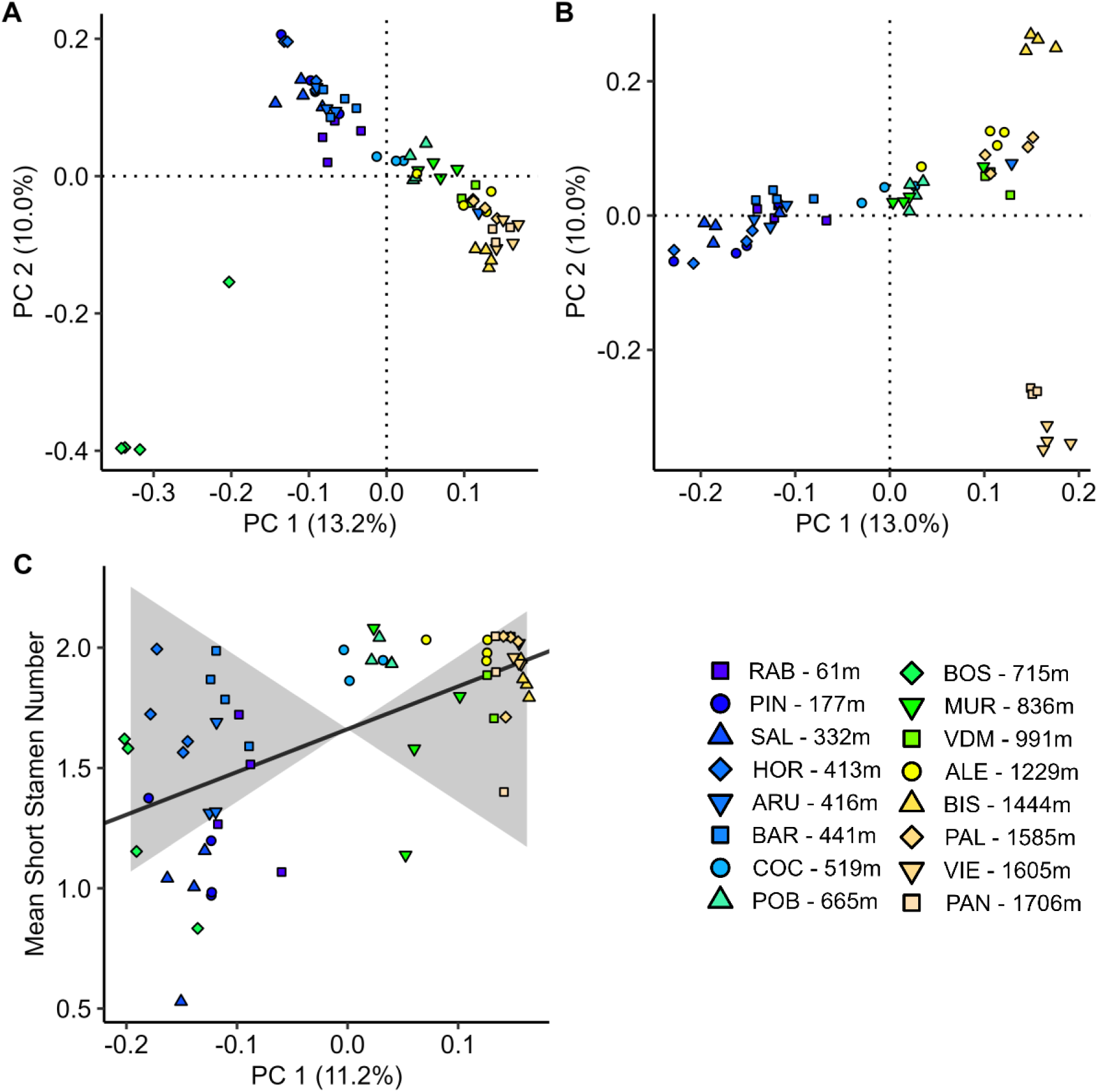
Genetic variation is correlated with elevation, but there is no evidence for divergent selection on short stamen number. **A)** Genetic PCA for PC1 and PC2 from 1,858,706 SNPs. **B)** Genetic PCA for PC1 and PC2 with BOS individuals excluded. **C)** Qpc results showing the relationship between mean short stamen number and the greatest axis of genetic differentiation, PC1, (black line, *p*=0.26) is within expectations due to neutral evolution (grey shading), as were PC2-4 (see text). The fill color of each point represents the population elevation. Panels A and C used slightly different input data (see Methods). **Alt text:** Graphs depicting genetic relatedness from genetic principal component analysis in panels A and B. In both panels, principal component 1 explains 13 percent of genetic variation and principal component 2 explains 10 percent. Panel C shows a positive relationship between mean short stamen number and principal component 1 but it is within expectations due to neutral evolution.

### Phenotyping

To assess genetic differentiation for short stamen number, plants were grown in growth-chamber common gardens in three blocks at Michigan State University, East Lansing, MI. Seeds were stratified at 4°C for five days before we increased temperatures to 22°C:18°C for 16-hour days at a constant 60% humidity for four weeks. After emergence, plants were thinned to one seedling per cell in a 200 cell plug tray. One to four plants from each of the 112 genotypes (median = 2; *N* = 230 plants) were vernalized at 4°C with 10-hour days for six weeks before returning to 22°C:18°C for 16-hour days through flowering.

On each plant, we counted stamen number (Fig. 1A) on up to three flowers at each of up to three timepoints throughout the duration of flowering to estimate short stamen number (1 to 9 flowers per plant; median = 6; *N* = 1,423 flowers). Short stamen number varied among flowers within all but three of the 58.2% of plants with stamen loss; the average standard error of short stamen number within individuals was 0.16. Population mean short stamen number was calculated as the arithmetic mean from all flowers phenotyped in a population (42 to 169 flowers per population; mean = 88.9; median = 99.5; average standard error within population = 0.06). The arithmetic population means were highly correlated with estimated marginal means for each population calculated with the *emmeans* package (Lenth, 2021) in R v4.2.2 (R Core Team, 2021) from a model with population as a fixed effect, random effects of flowering timepoint nested within plant nested within genotype which is nested within population, and random effect for block (*r* = 0.975; *p* < 0.001).

We sequenced the entire genomes of a subset of 61 genotypes (see below). The 61 genotypes include representatives from all 16 populations (3 or 4 different genotypes per population); we elected not to sequence all genotypes as prior work indicated low within-population variation (Gomaa et al., 2011; Montesinos et al., 2009). In the sequenced genotypes, stamen number was counted on up to four plants per genotype (median = 2; *N* = 141) and up to nine flowers per plant (median = 7; *N* = 971). For all analyses that incorporate both phenotypic and SNP information, arithmetic population means were recalculated as the mean from flowers scored for short stamen number from sequenced genotypes (18 to 118 flowers per population; median = 63). The population means from all flowers and the population means from only the sequenced genotypes are highly correlated (*r* = 0.985; *p* < 0.001) indicating the sequenced genotypes are representative of all plants scored for short stamen number.

### Testing for an elevational cline

We calculated both linear and quadratic regressions of short stamen number on elevation to identify if there is an elevational cline in short stamen number parallel to the latitudinal cline (Table S3). The quadratic regression was included because it increased the adjusted *R^2^* substantially even though the quadratic term was not significant.

### Sequencing

A subset of 61 genotypes (3-4 per population) were chosen for Ilumina paired-end whole-genome 150bp sequencing. Nextera adapter sequences were trimmed from the raw sequence data with Trim Galore (Krueger, 2019). We also clipped the first 15 bp of each read because quality checks with FastQC (Andrews, 2019) and MultiQC (Ewels et al., 2016) showed an identical, unusual, pattern in the first 10 bases of each read: GTTTTAAACT. Reads were mapped to the TAIR10 reference genome (Berardini et al., 2015) using BWA mem with default settings (Li & Durbin, 2009). The mean mapping rate for properly paired reads was 96.6% with a mean of 15,224,790 properly paired and mapped reads per genotype and a total of 943,936,960 mapped and paired reads in the dataset. The median depth across genotypes was low but acceptable at 8X. There is variation in coverage, missing data, and quality scores between genotypes, but no genotypes were excluded on this basis in order to maximize sample size (Table S2). Duplicate reads were marked with the Genome Analysis Toolkit (GATK) v4.1.4.1 (Van der Auwera & O’Connor, 2020) MarkDuplicates Spark. After an initial round of Haplocaller and GenotypeGVCF, the dataset was filtered with the parameters suggested by GATK best practices (Van der Auwera & O’Connor, 2020): QD <2, FS > 60, MQ < 40, ReadPosRankSum < −8, and MQRankSum < −12.5. The filtered file was used as a known variants file for base quality score recalibration (BQSR). Variants were then called again with HaplotypeCaller using the GVCF flag to keep all sites before combining samples for GenotypeGVCF with the all-sites flag to create a dataset that includes both variant and invariant sites (116,855,685 total sites).

### Testing the drift hypotheses

To estimate the effects of past drift and gene flow in each population, genome-wide pairwise nucleotide diversity (pi) was calculated for each population as an approximate measure of effective population size with pixy v1.0.4 (Korunes & Samuk, 2021) from a filtered dataset containing variant and invariant sites based on the pixy protocol. We used quality filters to remove all indels and low-quality sites that met the following criteria: quality score less than 20, mean depth less than 3, and more than 25% missing data. The filtered dataset contains 99,202,614 sites. A peak in nucleotide diversity was observed in the centromere of each chromosome, likely caused by fewer mapped sites (Korunes and Samuk, *pers. com.*). However, excluding previously published centromeres (Clark et al., 2007) did not meaningfully change genome-wide nucleotide diversity (*r* = 0.998; *p* <0.001), so centromeres are included in all of the following analyses (results excluding the centromeres can be found in Figs. S5-S7).

To test if variation in effective population size is contributing to a cline in short stamen number, we used multiple regression to identify how short stamen number was predicted by nucleotide diversity and elevation. While elevation is correlated with climatic variables such as precipitation and temperature (Montesinos-Navarro et al., 2011) that could be selective agents for plant traits, we used elevation because previous work in these same 16 populations demonstrated that elevation explained 54% of the variance in trait principal component 1 (traits include: phenology, water use efficiency, instantaneous CO_2_ and H_2_O exchange, PSII quantum efficiency, and specific leaf area) while the climate PC1 only explained 36% of the trait variation (Wolfe & Tonsor, 2014). We then visualized these results by using the residuals of single regression models. The visualized results underestimate the confidence interval because they do not account for uncertainty in calculating the residuals in the single regression models; the reported statistics are from the full model. We used *lme4* in R for all models (Bates et al., 2015).

A negative relationship between nucleotide diversity and short stamen number after correcting for elevation would support the hypothesis that smaller effective population sizes are constraining trait loss due to low standing variation and less effective selection. A relationship between short stamen number and elevation after correcting for nucleotide diversity could be evidence for divergent selection because it indicates the short stamen number cline persists beyond the variation explained by nucleotide diversity. However, a relationship between short stamen number and elevation could also result from shared evolutionary history, sometimes called shared genetic drift, if relatedness among populations is associated with elevation (Colautti & Lau, 2015). We cannot distinguish divergent selection from relatedness in the multiple regression analysis. Therefore, we further investigated evidence for divergent selection by testing for local adaptation in short stamen number.

### Testing for local adaptation

We used plink v1.9 (Chang et al., 2015) to filter the all-sites output from GATK to a “variant sites only” dataset for conducting genetic principal component analysis (PCA). The “variant sites only” dataset was also used for a genome-wide association study (GWAS; see below). We used quality filters to remove all indels and non-biallelic sites in addition to low quality sites that met the following criteria: minor allele frequency less than 5%, quality score less than 25, mean depth less than 5, and more than 25% missing data. The variant sites dataset contains 1,858,706 SNPs. Filtering parameters were chosen to maintain the same percent nonsynonymous sites, annotated by snpEff (Cingolani et al., 2012), as stricter parameters that removed sites with greater than 15% missing data and a quality score less than 60 while retaining more variants in the dataset. Genetic PCA was calculated for the first 20 PCs with plink v1.9 (Chang et al., 2015) to characterize genetic relatedness within, and differentiation between, populations. We looked for correlations between the population average PC value and elevation for each of the first 4 PCs to identify population structure associated with elevation.

We tested for divergent selection across populations on short stamen number, i.e., selection for more stamens at high elevation and/or fewer stamens at low elevation than expected due to genetic drift, with Qpc using the *quaint* R package (Josephs et al., 2019). Qpc is an extension of Qst-Fst analysis that uses genetic PCs to estimate additive genetic variance within and between populations (Josephs et al., 2019), and tests for phenotypic differentiation due to selection beyond that expected from neutral evolution. This differs from the multiple regression test for the drift hypothesis described above because Qpc explicitly considers among population differentiation. Qpc incorporates genetic relatedness within and among populations through a kinship matrix, rather than a single measure of population differentiation as in Qst-Fst. Qpc then tests for excess phenotypic divergence along major axes of genetic relatedness (principal components of the kinship matrix) rather than excess phenotypic divergence between populations as in Qst-Fst (Josephs et al., 2019). The input kinship matrix was generated from a random subset of 50,000 SNPs that had no missing data. This kinship matrix is slightly different from the kinship matrix used for genetic PCA because Qpc requires a kinship matrix standardized across all loci, not each locus individually as in plink, and *quaint* is not capable of dealing with missing data.

### Using genome wide association studies (GWAS) to identify loci associated with short stamen loss

We performed GWAS to find genomic regions associated with short stamen number. Mean short stamen number was not normally distributed (Shapiro-Wilk *w* = 0.85; *p* = 3.27×10^-6^; Fig. S1). Arcsine transformation made the distribution closer to normal but still skewed (Shapiro-Wilk *w* = 0.91; *p* = 2.72×10^-4^; Fig. S1). Phenotypic and genotypic data were merged in plink v1.9 (Chang et al., 2015). GWAS was conducted in gemma v0.98.4 with the Wald hypothesis test for each SNP against the alternate allele and a centered kinship matrix to account for population structure (Zhou & Stephens, 2012). The output was visualized with the *qqman* (Turner, 2018) and *ggplot2* (Wickham et al., 2019) R packages. SNPs were further investigated if they passed a significance threshold specified using a false discovery rate (FDR) < 0.05 determined with p.adjust in R.

Because the arcsine-transformed stamen number still showed some skew, we conducted GWAS using additional strategies for quantifying short stamen number. We ran a GWAS with untransformed mean short stamen number and another with mean short stamen number coded as a binary trait. In the latter, genotypes with no short stamen loss (mean short stamen number = 2) were coded as controls and genotypes with any amount of short stamen loss (mean short stamen number < 2) were coded as cases. Finally, a fourth GWAS was conducted on the subset of genotypes that experience short stamen loss (i.e., GWAS on only the genotypes with mean short stamen number < 2; Shapiro-Wilk *w* = 0.93; *p* = 0.0132) to ameliorate the zero-inflated-like distribution in the other continuous GWAS caused by excess genotypes with a mean short stamen number equal to 2 (Fig. S1). In total we ran four GWAS on different ways of quantifying short stamen number: arcsine transformed, untransformed, binary coded, and the subset with short stamen number less than two.

We identified genetic regions with SNPs associated with short stamen number in at least two GWAS to find candidate regions for further study (“shared” SNPs). We identified overlapping regions associated with stamen loss among the four GWAS analyses by creating a 1kb window centered on each SNP that had an FDR adjusted p-value below 0.10 in any GWAS and searching for SNPs within that window that were also below an FDR adjusted p-value of 0.10 in at least one other GWAS. We chose a more lenient FDR of 0.10 for this analysis because we have higher confidence SNPs are associated with short stamen number if they pass a significance threshold in multiple analyses. We chose a window of 1kb to include regions flanking the associated SNP, although we recognize that *A. thaliana* genes can be larger than 1kb and that linkage disequilibrium (LD) begins to drop off around 50kb in our individuals (Fig. S2), so this is a conservative overlap criterion. SNPs in overlapping stamen loss associated regions were investigated with the TAIR10 genome browser. All figures were created with *ggplot2* (Wickham et al., 2019) and *ggpubr* (Kassambara, 2023) unless otherwise noted.

## Results

### Short stamen loss is more common at low elevation

Mean short stamen number, when measured in a common garden, increases with the elevation of the source population until reaching close to two stamens in plants from ∼1300m (quadratic regression: β = 4.84×10^-4^; *p* = 0.004; γ = −4.73×10^-7^; *p* = 0.137; Fig. 2A; Table S3). Thus, short stamen retention is more common in populations from higher elevations and loss is more common at low elevations in northeast Spain. This elevational cline is similar to the previously identified latitudinal cline where short stamen loss is more common at southern latitudes (Royer et al., 2016).

### Nucleotide diversity decreases with elevation, but there is no evidence that variation in effective population size contributes to the cline in short stamen number

Consistent with expectations that *A. thaliana* in the Pyrenees experienced repeated founder effects after the last glaciation, high elevation populations have less nucleotide diversity than low elevation populations (β = −1.81×10^-6^; *p* < 0.001; Fig. 2B; Table S3). The three populations with the lowest nucleotide diversity are high elevation populations BIS, VIE, and PAN. The results are consistent with prior findings that genetic diversity and effective population size decrease with elevation in this region (Gomaa et al., 2011; Montesinos et al., 2009). Population nucleotide diversity is comparable to nucleotide diversity of *A. thaliana* across the Iberian Peninsula and the European range (The 1001 Genomes Consortium, 2016).

We hypothesized a negative relationship between nucleotide diversity and short stamen number if variation in effective population size was contributing to the short stamen cline, but nucleotide diversity did not significantly predict short stamen number when accounting for elevation (β = - 138.5; *p* =0.167; Fig. 2D; Table S3). Further, the relationship between short stamen number and elevation is similar whether nucleotide diversity is included or not (Fig. 2C, compare to Fig. 2A; Table S3). These results suggest that variation in effective population size is not contributing significantly to the short stamen number cline. Rather, local adaptation, correlated responses to selection, or shared evolutionary history may be contributing to the cline.

### There is no evidence for divergent selection on short stamen number along the elevational cline

An elevational cline in short stamen number could result from local adaptation to different optima at high and low elevations. To demonstrate evidence of local adaptation resulting from divergent selection, we would need to show that the cline in short stamen number is stronger than what could be generated by neutral evolution alone. Genetic structure, here measured using genetic PCA, shows a strong pattern of genetic differentiation by elevation; both PC1 (*r* = −0.75; *p* = 9.22×10^-4^) and PC2 (*r* = −0.54; *p* = 0.03) are correlated with elevation (Fig. 3A). Neither PC3 nor PC4 are correlated with elevation (Fig. S3). The BOS population is an outlier along both PC1 and PC2 (Fig. 3A). This aligns with geographic information because BOS is located on a northern face of the Spanish Pyrenees while the other populations are on southern faces, and prior work identified BOS in the Northwestern genetic cluster of the Iberian Peninsula while the other 15 populations belong to the Northeast cluster or are classified as mixed (Castilla et al., 2020). The PCA was conducted a second time after removing the BOS population (Fig. 3B). These results continue to show an elevational cline along PC1 (*r* = −0.92; *p* = 1.15×10^-6^), though PC2 (*r* = −0.17; *p* = 0.55) separates the four highest elevation populations from each other with the closest clustering of PAN and VIE. As before, neither PC3 nor PC4 are correlated with elevation (Fig. S3).

The strong genetic differentiation across elevation we observed means that we have low power to test for even greater differentiation in stamen loss as evidence for divergent selection. Thus it is not very surprising that we did not find evidence for divergent selection on short stamen number with Qpc. Differences in short stamen number between high and low elevation populations were not larger than could be explained by neutral evolution in any of the first four PCs, which together explain 40.2% of the genetic variation (Fig. 3C; *p* = 0.257, 0.914, 0.802, 0.795 in order). Therefore, the Qpc analysis does not support local adaptation contributing to the cline, but we cannot definitively rule out local adaptation because of the low power of our analysis.

### Parallel clines in short stamen have different genetic bases

We conducted GWAS to identify loci associated with short stamen number. Because short stamen number was not normally distributed (Fig. S1), we conducted GWAS with four strategies for quantifying short stamen number: arcsine-transformed mean short stamen number, untransformed mean short stamen number, a binary trait of whether or not individuals lacked any short stamens, and an untransformed subset of only genotypes that lack some short stamens. Each GWAS found different SNPs associated with the trait. The qqplot for the arcsine-transformed GWAS shows most p-values close to the 1:1 line between Expected and Observed (Fig. 4B) but the p-values in the binary GWAS deviate from the 1:1 line (Fig. S4B), indicating a high false positive rate in identifying SNPS associated with any short stamen loss. In accordance with this observation, the binary GWAS had 1,707 SNPs associated with short stamen loss at FDR < 0.05 while the other analyses only had two or three (Figs. 4, S4). Only two SNPs are associated with arcsine-transformed short stamen number (FDR < 0.05; Fig. 4). The SNP at Chr3:2942726 is within a FAD/NAD(P)-binding oxidoreductase family protein (AT3G09580) that localizes in the chloroplast (Tomizioli et al., 2014). The SNP at Chr5:13458838 is not within a gene; the closest gene, *PICALM3* (AT5G35200), is just over 3kb away.

**Figure 4.**
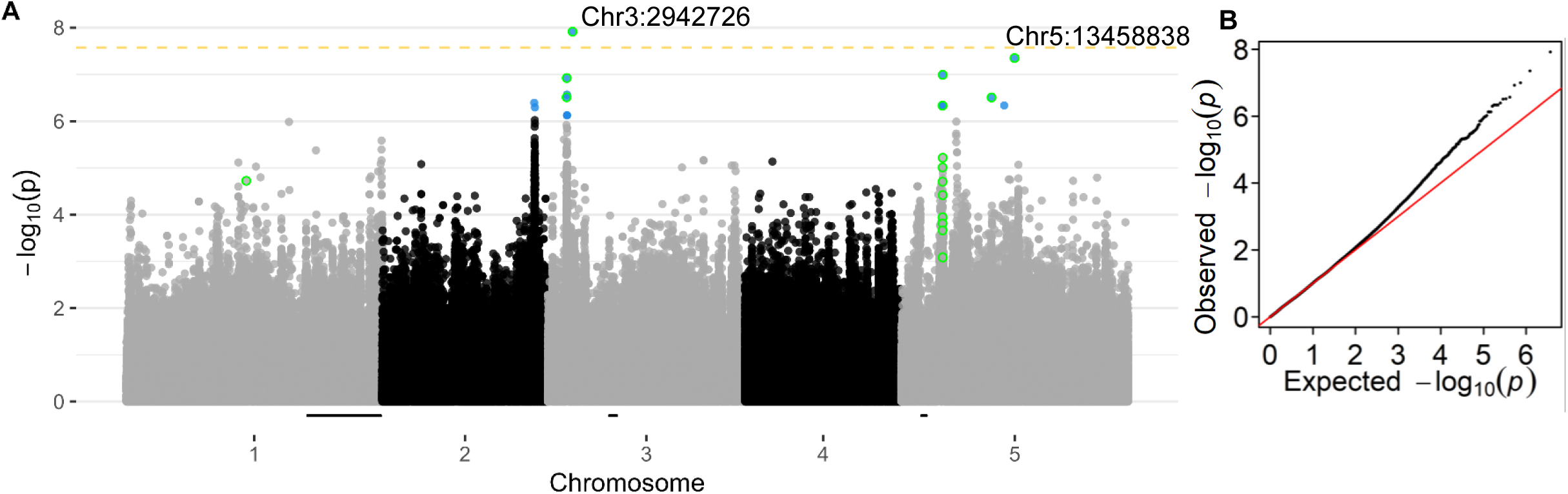
Few SNPs are associated with short stamen number. Manhattan plot **(A)** and QQ plot **(B)** for arcsine-transformed mean short stamen number. The yellow dashed line indicates significance at p=0.05 after Bonferroni correction. Blue points are significant below a FDR of 0.10. Points labeled with their chromosomal location are significant below a FDR of 0.05. Points with a green outline are shared between at least two short stamen GWAS below FDR 0.10 (n = 20). The black bars on the x axis are Bayes 95% credible intervals for short stamen number QTL identified by Royer et al. (2016). **Alt text:** Graphs showing results of arcsine-transformed genome wide association study. The two labeled points in panel A are at chromosome 3 position 2942726 and on chromosome 5 at 13458838. There are eight points significant below a false discovery rate of 0.10, most of them are outlined in green. Half of the shared SNPs are not significant in this GWAS.

We identified top candidates for loci associated with short stamen number by identifying 1kb windows that have SNPs associated with short stamen number in multiple GWAS (“shared” SNPs). To do so, we calculated 1kb windows centered on all SNPs associated with short stamen number (FDR < 0.10) and searched for other SNPs that fit the same criteria and fell within each window. Twenty SNPs were associated with short stamen number in more than one GWAS from 9 different 1kb windows (Table S4). None of the SNPs fall within genes with known stamen function. However, a shared SNP at Chr3:2253161 is approximately 40kb away, thus within LD (Fig. S2), from *FHA2* (AT3G07220), a SMAD/FHA domain containing protein involved in stamen development (Ahn et al., 2013; Gu et al., 2020). Mutating *FHA2* proteins can cause plants to have fewer stamens, though flowers sometimes lose long stamens and sometimes lose short stamens (Ahn et al., 2013). The two SNPs associated with arcsine-transformed short stamen number (FDR < 0.05) are included in the shared SNPs (Fig. 4; Table S4).

All of the SNPs associated with variation in short stamen number fall outside the 95% credible intervals of the previously identified QTL for short stamen number that used recombinant inbred lines with parents from the extremes of the latitudinal short stamen loss gradient (Royer et al., 2016). The closest intersection is on chromosome 5, where SNPs associated with short stamens are approximately 2,000kb further into the chromosome than the previously identified short stamen loss QTL. This is beyond the start of LD decay we estimated of 50kb (Fig. S2). Thus, the parallel phenotypic clines in short stamen number across elevation and latitude are not parallel genetically.

## Discussion

In this study, we tested for an elevation cline in short stamen loss. We found that short stamen number increased with elevation (Fig. 2A), similar to the cline of more short stamens at higher latitudes observed by Royer et al. (2016). Short stamen number is one of many traits that show an elevational cline in this region (Montesinos et al., 2009; Montesinos-Navarro et al., 2011; Wolfe & Tonsor, 2014). Of these traits, days to bolting shows similar elevational and latitudinal clines with delayed bolting at high elevations and northern latitudes (Montesinos-Navarro et al., 2011; Stinchcombe et al., 2004). Thus, short stamen number joins a growing number of traits with an elevational cline in northeast Spain and a much smaller set of traits with parallel clines in elevation and latitude. These clines could be caused by effective population sizes because both high elevation populations and northern latitude populations may have undergone repeated bottlenecks that reduced effective population size since the last glaciation (Beck et al., 2008; François et al., 2008; Hughes et al., 2006; Lewandowska-Sabat et al., 2010; Oakley et al., 2019). While northeast Spain does include putative glacial refugia regions (Médail & Diadema, 2009), none of the 16 populations analyzed here have been classified as relict lineages that predate the last glaciation (Castilla et al., 2020). Selection could also contribute to similar latitudinal and elevational clines because environmental factors such as temperature often vary in the same way across latitude and elevation.

We then tested if the cline in short stamen loss was caused by variation in effective population size if a combination of low genetic variation and less effective selection in high elevation populations hinders a response to selection against short stamens. We found variation in genetic diversity across elevations consistent with expectations from repeated founder effects during range expansion after the last glaciation and/or low gene flow (Fig. 2B). However, the multiple regression suggests that variation in genetic diversity is not contributing to the short stamen number cline (Fig. 2C and 2D). Instead, the multiple regression suggests that elevation can explain both variation in genetic diversity and in stamen number. It is important to note that our phenotypic clines show populations with low genetic diversity retain two short stamens, consistent with our hypothesis. However, studies that identify a role of genetic drift in trait loss have hypothesized the opposite, where populations with low genetic diversity have trait loss caused by drift (Eckert et al., 1996; Lahti et al., 2009).

Next, we tested if local adaptation was contributing to the cline in short stamen loss using Qpc, an extension of Qst-Fst (Josephs et al., 2019). We found no evidence of local adaptation contributing to the cline in short stamen number (Fig. 3C). One caveat to our findings is that Qst-Fst struggles to identify selection when there is large genetic differentiation between populations or weak selection that results in similar values for Qst and Fst (Whitlock & Guillaume, 2009).

This is also potentially an issue for Qpc in this study as there is a great deal of genetic differentiation between high and low elevation populations (Fig. 3A). However, the Qst-Fst approach has identified local adaptation contributing to elevational clines in leaf succulence and specific leaf area in 14 populations of *A. thaliana* from the Swiss Alps where there is also a great deal of genetic differentiation (Luo et al., 2015). Qpc has also previously been used to identify local adaptation in 249 natural *A. thaliana* accessions from across the European native range; local adaptation was identified in initial size, growth rate at 16°C, and temperature response (Clauw et al., 2022), so there is still potential to use Qpc to identify local adaptation in systems with strong genetic differentiation. Ultimately, reciprocal transplant studies across elevations in the field will provide the strongest evidence for or against local adaptation, but genomic approaches like Qpc will still be necessary when reciprocal transplants are not possible or when many populations are being studied as we do here.

Finally, we used GWAS to identify loci associated with short stamen number. We identified 20 SNPs associated with short stamen number in multiple GWAS (FDR < 0.10). These SNPs lie outside the QTL for short stamens that were previously identified in a cross between a northern European and southern European accession (Royer et al., 2016). The untransformed effect sizes of our loci (Table S4) range from an absolute value of 0.23 to 0.33 and are much larger than the effect sizes of the QTL identified in Royer et al. (2016), untransformed effect sizes range from 0.05 to 0.17, suggesting that we may be missing the previously identified QTL due to lower power in the GWAS. Alternatively, the populations studied here may be fixed for the southern allele identified in Royer et al (2016). However, our GWAS effect sizes may be inflated due to winner’s curse (Göring et al., 2001; Josephs et al., 2017) which causes effect sizes to be overestimated in studies with small sample sizes such as ours (Capen et al., 1971). In addition, the QTL identified in Royer et al. (2016) were epistatic, which could make them harder to detect through standard GWAS approaches. Further, many of the loci underlying short stamen loss likely have effect sizes too small to detect by the methods we used and by the previous QTL analysis. For example, a QTL study in maize identified 50 QTL associated with a trait they artificially selected for, yet these QTL explained only half the genetic divergence for the trait indicating many other QTL were involved but not detected (Laurie et al., 2004). Overall, our results suggest that the latitudinal cline in short stamen number across Europe has a different genetic basis than the elevational cline in short stamen number observed in northeast Spain.

While the short stamen loss clines are similar between latitude and elevation, the lack of overlap between the associated loci suggests that short stamen loss is caused by variants at different genes in different geographic regions. These results are consistent with other findings that the chromosome regions underlying the same phenotypic cline can differ by region. For example, elevational clines of *A. thaliana* in different geographic regions show delayed flowering at higher elevations but the genetic basis of this cline varies by global region (Gamba et al., 2024). Additionally, parallel clines in flowering time in *A. thaliana* in the Cape Verde Islands arose from mutations at different genes, *FRI* and *FLC* (Fulgione et al., 2022). Even in outcrossing *Arabidopsis* species, functional parallel evolution is more common than genetic parallel evolution (Bohutínská et al., 2021). The lack of genetic parallelism is not surprising but may complicate future work to identify the developmental pathways resulting in loss of short stamens.

In conclusion, short stamen loss has occurred in low elevation populations more than in high elevation populations. Our results suggest that retention of two short stamens in high elevation populations is not explained by reduced effective population sizes or by local adaptation to different optima in high and low elevation populations. Thus, the evolutionary mechanisms underlying variation in short stamen number are excitingly complex and deserve further study. A third hypothesis is that correlated responses to selection may contribute to the cline in short stamen number (Futuyma, 2010; Lande, 1979). However, correlated responses to selection on other locally adapted traits could result in weak evidence for divergent selection; we did not find this. Future work should better characterize direct selection and correlated responses to selection on short stamens in the field by measuring fitness of plants with natural variation in short stamen number and experimental manipulation of stamens, like that done by Royer et al. (2016), in the field. Further, refining the short stamen number QTL to candidate genes and comparing to genetic regions underlying locally adapted traits could uncover the role of linked or pleiotropic genes leading to correlated responses to selection. These additional studies across both latitudinal and elevational gradients will characterize the interplay of direct selection, correlated responses to selection, and genetic drift in trait loss and identify parallel evolutionary forces at play across environmental contexts.

## Author Contributions

SGP collected the phenotypic data and conceived the research with JKC. JRP oversaw the sequencing. AG and GSB did preliminary analyses and contributed to conceptual development. CGO, SJT and FXP provided seed material and guidance in project development. SFB analyzed the data and drafted the manuscript with guidance from EBJ and JKC. JKC managed the project. All authors provided comments on the manuscript.

## Funding

Research reported was supported by the National Institute of General Medical Sciences of the National Institutes of Health (NIH) under award R35GM137919 (awarded to GSB) and R35GM142829 (awarded to EBJ), and by National Science Foundation awards DEB 0919452 (to JKC) and DEB 2223962 (to JKC and GSB). The content is solely the responsibility of the authors and does not necessarily represent the official views of the NIH or NSF.

## Data Availability

These sequence data will be submitted to the GenBank database upon manuscript acceptance. All phenotypic data and code will be made available on GitHub upon acceptance. GWAS output files will be made available on Dryad upon acceptance.

## Conflict of Interest

The authors declare no conflicts of interest.

## Acknowledgements

We thank D. Schemske for guidance throughout the initial stages of the project, N. Zhang for library preparation, and A. Platts for bioinformatics assistance. This manuscript was improved by comments from Conner lab and Josephs lab members. This work is not possible without the foundational work of Dr. Steven Tonsor; all authors agree on his inclusion in this manuscript and JKC accepts all responsibility for his contribution. We dedicate this paper to his memory.

## Notes

### Competing Interest Statement

The authors have declared no competing interest.

### Summary of Updates

Introduction and Discussed updated to clarify hypotheses; Methods edited for clarity; Figure 1 revised

