## Supplemental Information for "Evaluating the Roles of Drift and Selection in Trait Loss along an Elevational Gradient"

**Supplemental Figures:**

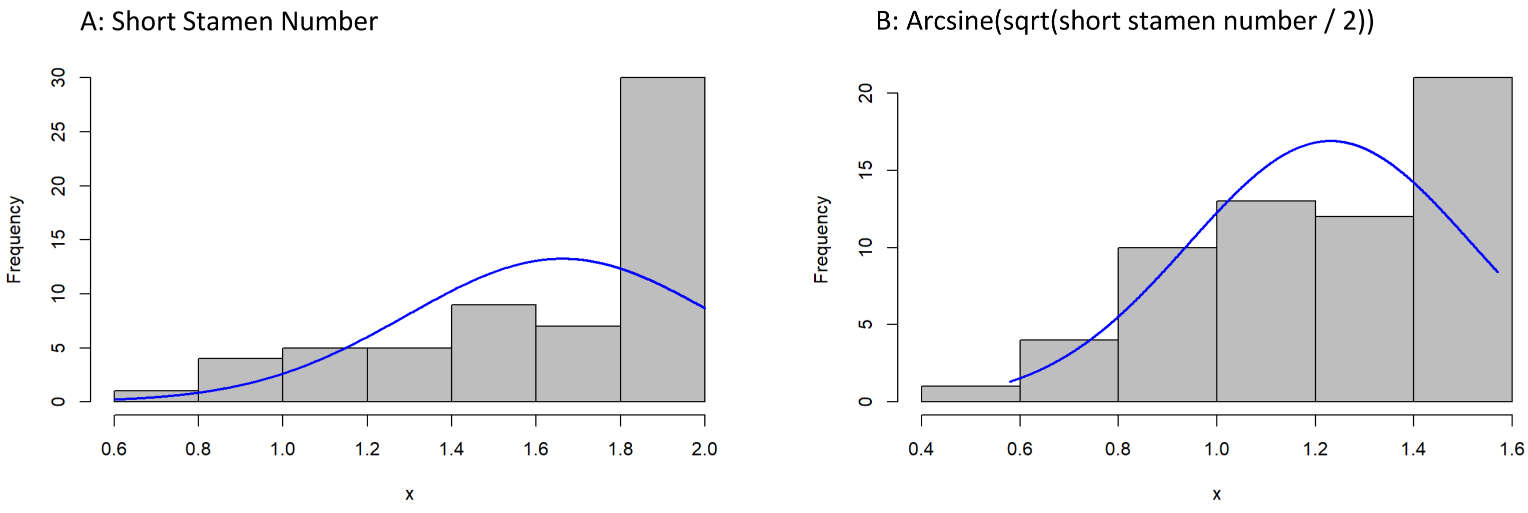

**Supplemental Figure 1. Short stamen number is not normally distributed, even after transformation.** Histograms of individual mean short stamen number without transformation (A) and after arcsine transformation (B).

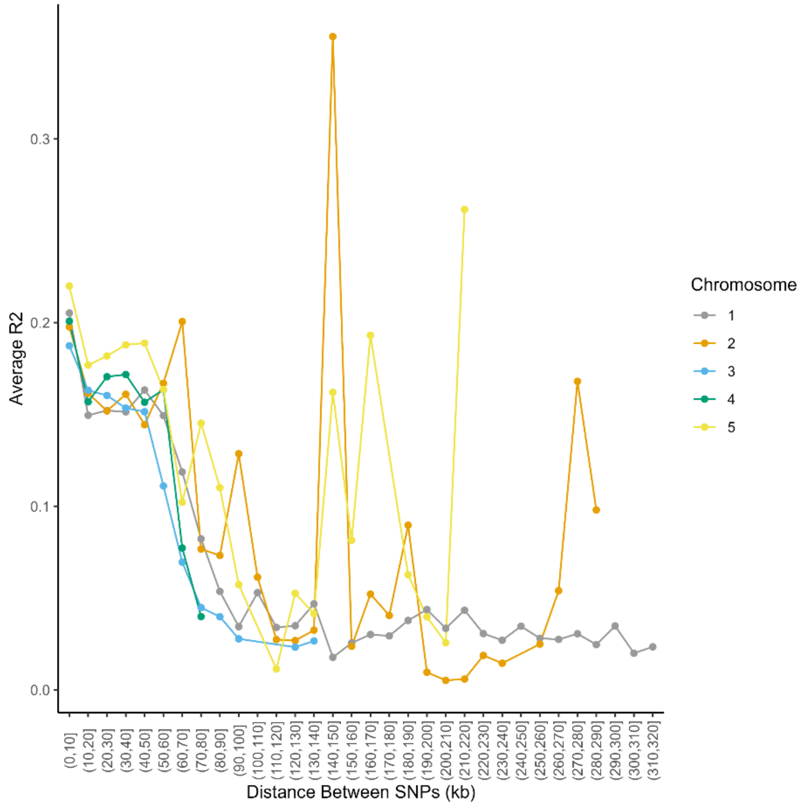

**Supplemental Figure 2. Linkage Disequilibrium decay within each chromosome.** LD decay calculated with plink. We estimated the first major drop in LD for all chromosomes around 50kb. X axis is 10kb windows from 0 to 320kb to bin the distance between SNPs.

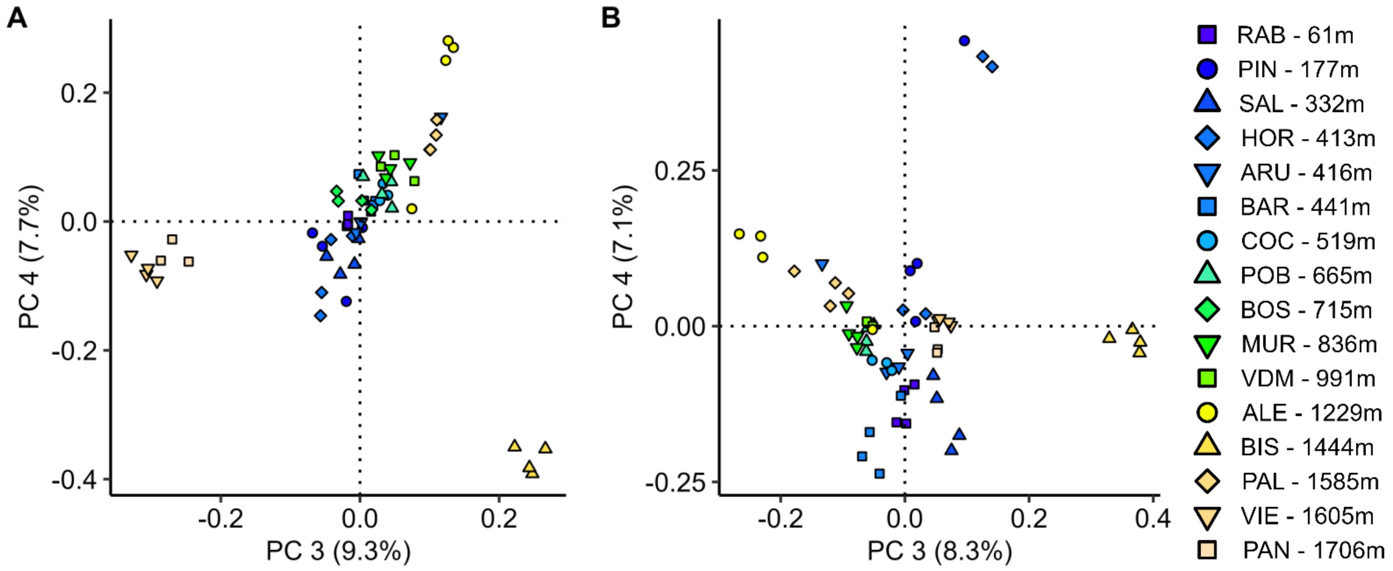

**Supplemental Figure 3. PC3 and PC4 are not correlated with elevation.** Genetic principal component analysis for PC3 and PC4 with all populations (A) and after excluding all BOS population individuals (B). With all populations, PC3 (r = -0.17, p = 0.52) and PC4 (r = -0.08, p = 0.74) are not correlated with elevation. Similarly, with BOS excluded neither PC3 (r = 0.11, p = 0.69) nor PC4 (r = 0.14, p = 0.61) are correlated with elevation.

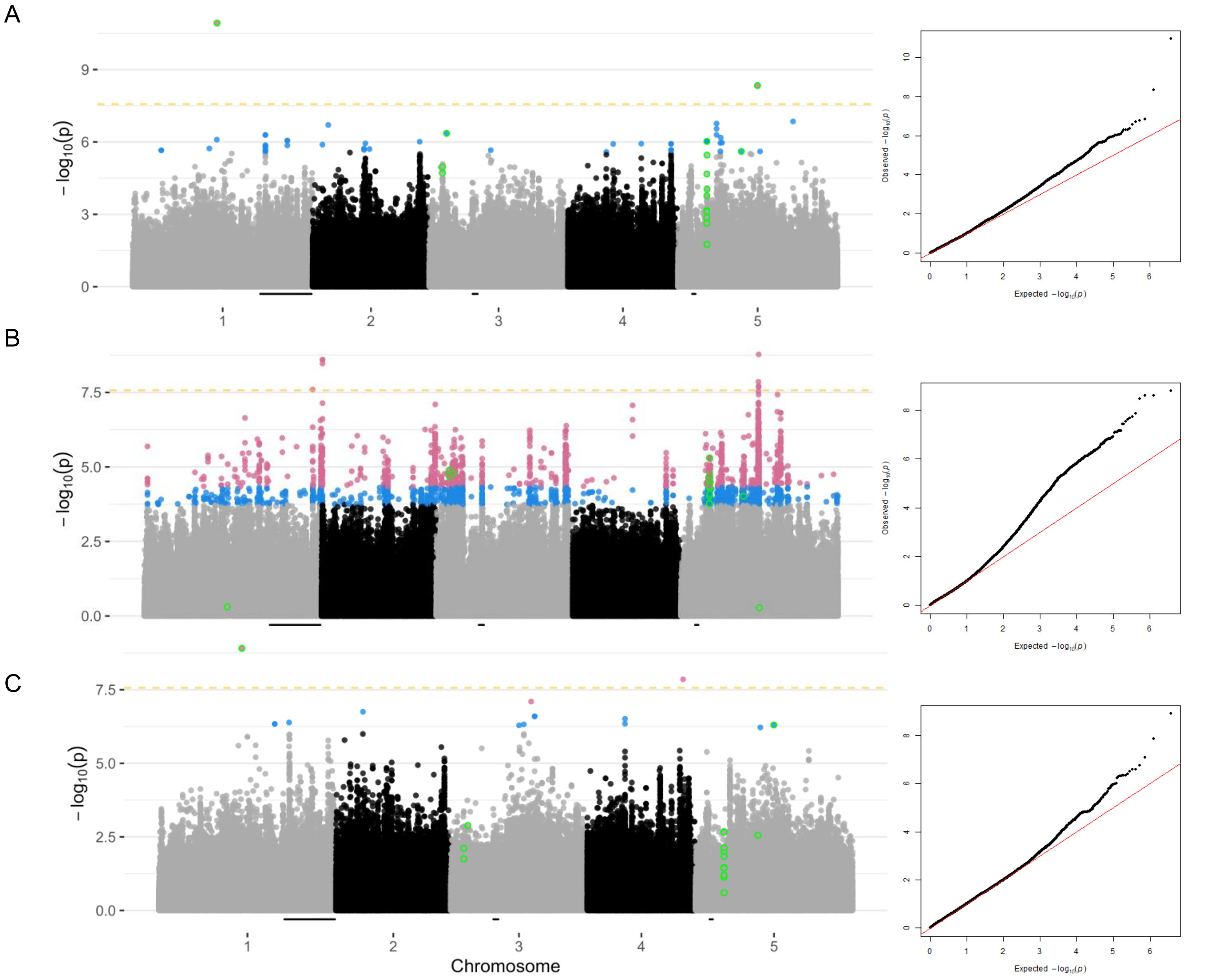

**Supplemental Figure 4. Additional manhattan plots for GWAS for short stamen number.** (A) Manhattan plot for untransformed mean short stamen number. (B) Manhattan plot for short stamen number coded as a binary trait (mean short stamen number < 2 coded as cases). (C) Manhattan plot for only the subset of individuals that showed any short stamen loss (if mean short stamen number < 2, individual included in this analysis). The yellow dashed line indicates significance at p=0.05 after Bonferroni correction. Blue points are significant below a FDR of 0.10. Pink points are significant below a FDR of 0.05. Points with a green outline are shared between at least two GWAS below a FDR 0.10 (n = 20). The black bars on the x axis are Bayes 95% credible intervals for short stamen number QTL identified by Royer et al. (2016).

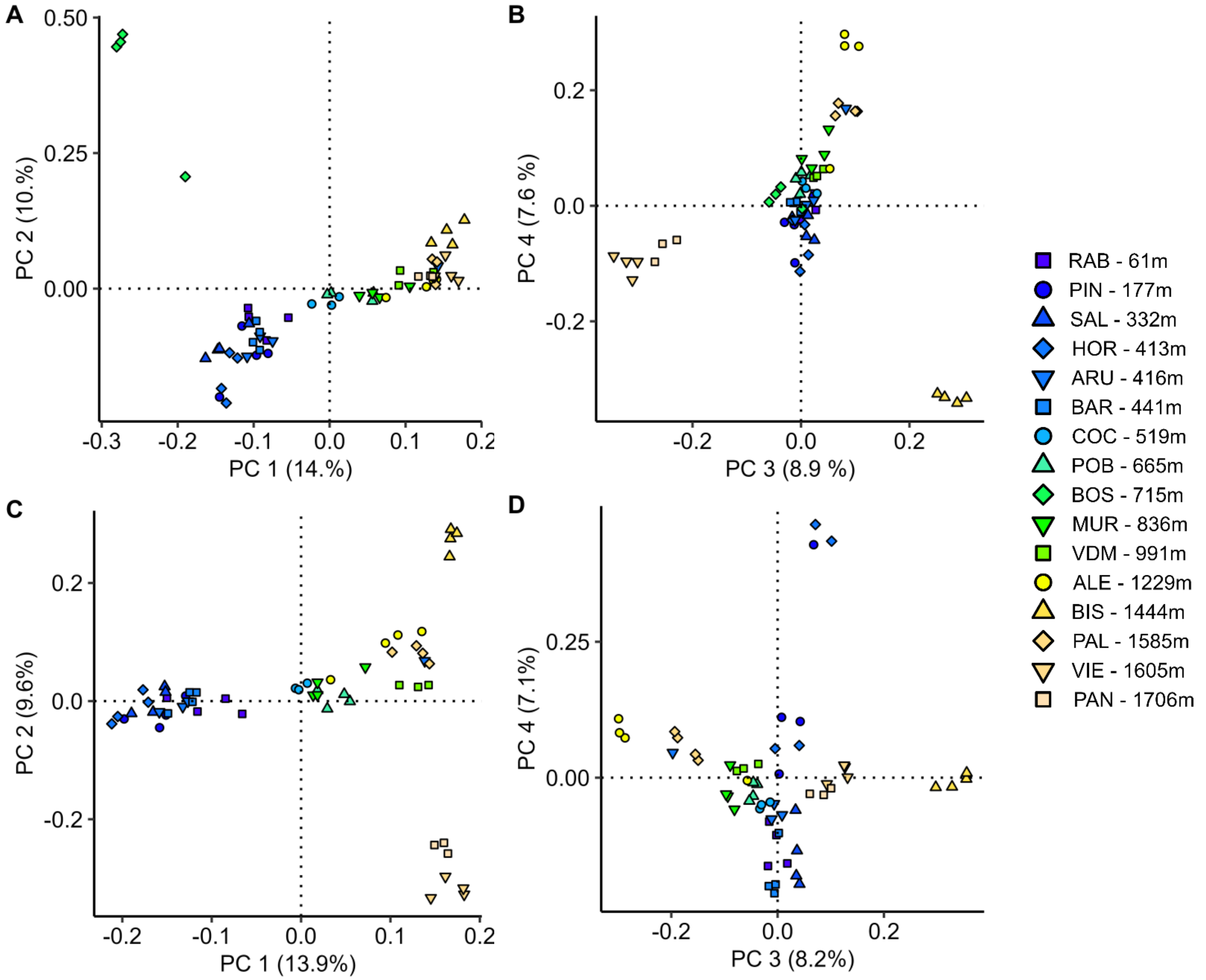

**Supplemental Figure 5. Genetic Principal Component Analysis without the centromere region.** Patterns of relatedness from genetic principal component analysis do not meaningfully change after excluding the centromere whether the BOS population is included (A, B) or excluded (C, D).

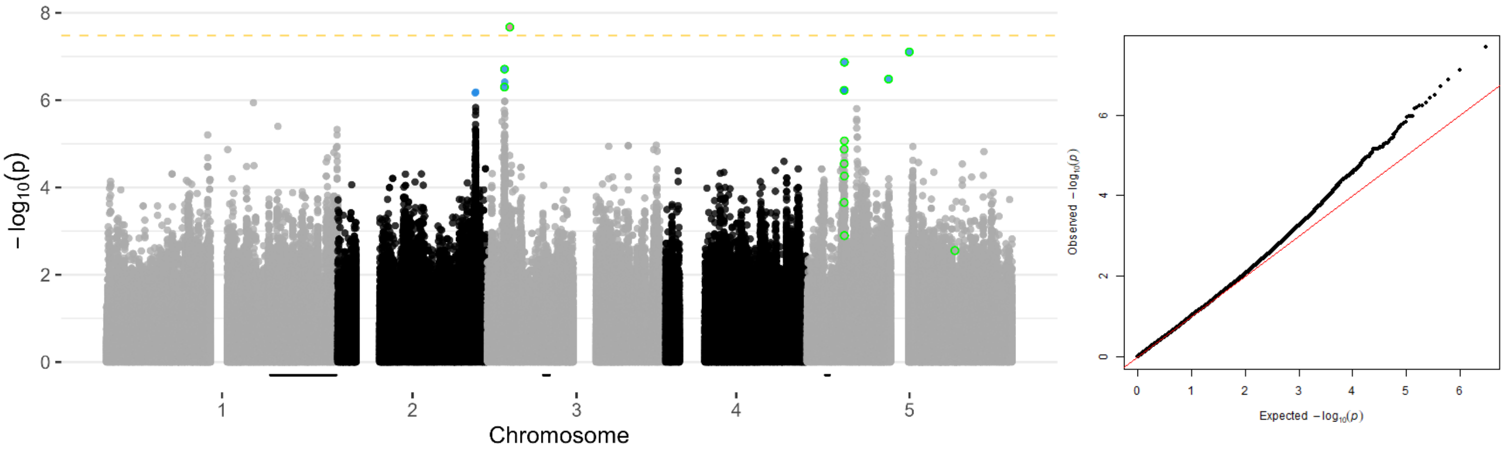

**Supplemental Figure 6. Manhattan plot for short stamen number after excluding the centromere region.** Manhattan plot for arcsine transformed mean short stamen number. The yellow dashed line indicates significance at p=0.05 after Bonferroni correction. Blue points are significant below a FDR of 0.10. Pink points are significant below a FDR of 0.05. Points with a green outline are shared between at least two GWAS below a FDR 0.10 (n = 16). The black bars on the x axis are Bayes 95% credible intervals for short stamen number QTL identified by Royer et al. (2016).

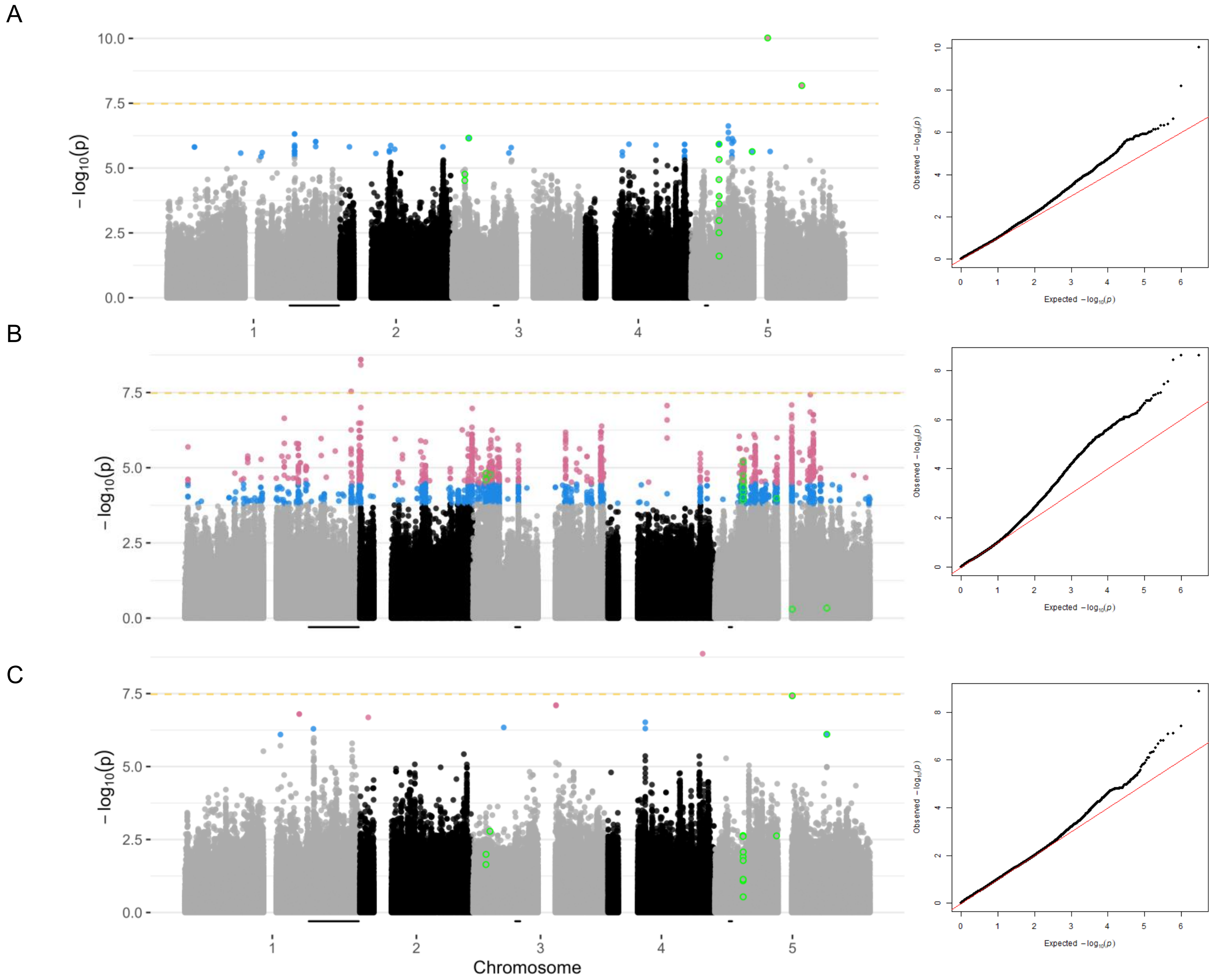

**Supplemental Figure 7. Additional manhattan plots for GWAS for short stamen number with the centromere excluded.** (A) Manhattan plot for untransformed mean short stamen number. (B) Manhattan plot for short stamen number coded as a binary trait (mean short stamen number < 2 coded as cases). (C) Manhattan plot for only the subset of individuals that showed any short stamen loss (if mean short stamen number < 2, individual included in this analysis). The yellow dashed line indicates significance at p=0.05 after Bonferroni correction. Blue points are significant below a FDR of 0.10. Pink points are significant below a FDR of 0.05. Points with a green outline are shared between at least two GWAS below a FDR 0.10 (n = 16). The black bars on the x axis are Bayes 95% credible intervals for short stamen number QTL identified by Royer et al. (2016).

**Supplemental Tables:**

**Table S1. Population metadata**. Population indicates the closest town to seed collection location. PopCode is a 3 letter abbreviation for each population. Elevation, latitude, and longitude data of seed collection site.

| Population | PopCode | Elevation (m) | Latitude (Decimal Degree) | Longitude (Decimal Degree) |
|---|---|---:|---:|---:|
| Rabos | RAB | 61 | 42.37 | 3.03 |
| Pineda de Mar | PIN | 177 | 41.66 | 2.66 |
| Santa Pellaia | SAL | 332 | 41.93 | 2.92 |
| Hortsavinya | HOR | 413 | 41.66 | 2.62 |
| Arbucies | ARU | 416 | 41.81 | 2.49 |
| Barcelona | BAR | 441 | 41.43 | 2.13 |
| Cap de Creus | COC | 519 | 42.32 | 3.16 |
| Poblet | POB | 665 | 41.35 | 1.03 |
| Bossost | BOS | 715 | 42.78 | 0.69 |
| Mura | MUR | 836 | 41.67 | 2 |
| Vilanova de Meia | VDM | 991 | 42.03 | 1.03 |
| Albet | ALE | 1229 | 42.41 | 1.32 |
| Bisaurri | BIS | 1444 | 42.49 | 0.53 |
| Pallerols de Canto | PAL | 1585 | 42.34 | 1.3 |
| Vielha | VIE | 1605 | 42.63 | 0.76 |
| Panticosa | PAN | 1706 | 42.76 | -0.23 |

**Table S2. Sequencing Quality.** For each line, total reads, the number of mapped reads, and the mapping rate from bwa are reported. The frequency of missing sites was calculated with vcftools after Base Quality Score Recalibration. Bamtools was used to report the percent of the genome with 1x and 4x coverage as well as median depth. Average quality score was extracted from vcf files with GATK. Mean, Median, and sum, where appropriate, are reported at the bottom of the table.

| Line | Total Reads | Mapped Reads paired | Mapping Rate paired | Frequency Missing Sites | Percent 1X | Percent 4X | Median Depth | Average Quality (NA excluded) |
|---|---|---|---|---|---|---|---|---|
| **ALE10** | 19712121 | 19184226 | 97.62 | 0.095284 | 0.93380832 | 0.848668447 | 9 | 26.710325 |
| **ALE12** | 19417011 | 18856422 | 97.48 | 0.0885923 | 0.938827568 | 0.89521716 | 10 | 30.779432 |
| **ALE16** | 3833518 | 3685146 | 96.39 | 0.358327 | 0.713647853 | 0.157023251 | 1 | 7.887054 |
| **ALE4** | 27137190 | 26132148 | 96.73 | 0.0779572 | 0.945231595 | 0.926439603 | 17 | 46.274956 |
| **ARB10** | 13753206 | 13249968 | 96.58 | 0.136072 | 0.918336322 | 0.728437677 | 6 | 18.480365 |
| **ARB3** | 21363640 | 20581102 | 96.62 | 0.0863734 | 0.94250462 | 0.870279153 | 10 | 30.108697 |
| **ARB6** | 20588486 | 20085288 | 97.8 | 0.101864 | 0.940417965 | 0.86537087 | 9 | 25.454609 |
| **ARB8** | 15833079 | 15240030 | 96.59 | 0.086599 | 0.93972272 | 0.861390169 | 9 | 26.609563 |
| **BAR11** | 7192507 | 6683048 | 93.04 | 0.380374 | 0.768546653 | 0.307549973 | 2 | 9.927844 |
| **BAR3** | 13209944 | 12849326 | 97.6 | 0.107214 | 0.933722864 | 0.825450573 | 8 | 23.911895 |
| **BAR4** | 16372218 | 15852820 | 97.17 | 0.081261 | 0.94403433 | 0.862889773 | 8 | 24.366533 |
| **BAR9** | 14955682 | 14190512 | 95.06 | 0.16471 | 0.906448036 | 0.66494117 | 5 | 17.04758 |
| **BIS11** | 8950013 | 8515326 | 95.34 | 0.246881 | 0.842364527 | 0.442343408 | 3 | 12.92224 |
| **BIS16** | 11657278 | 11174536 | 96.23 | 0.0878037 | 0.93895045 | 0.813676452 | 7 | 21.496708 |
| **BIS20** | 12101273 | 11628344 | 96.45 | 0.08588 | 0.941025012 | 0.811626785 | 7 | 20.769619 |
| **BIS8** | 13098954 | 12523234 | 95.86 | 0.131212 | 0.918908352 | 0.711081995 | 6 | 18.983814 |
| **BOS10** | 27285715 | 26509336 | 97.46 | 0.0757308 | 0.949908511 | 0.92948089 | 15 | 40.5501 |
| **BOS5** | 17913901 | 17379876 | 97.3 | 0.0827965 | 0.944301906 | 0.861507097 | 9 | 26.492931 |
| **BOS6** | 14892443 | 14533824 | 97.9 | 0.0875408 | 0.941322836 | 0.845040798 | 8 | 23.916785 |
| **BOS9** | 20317221 | 19721584 | 97.41 | 0.0739174 | 0.94939497 | 0.920368364 | 12 | 33.818551 |
| **COC14** | 19730554 | 19076102 | 96.97 | 0.0898506 | 0.941173031 | 0.865796868 | 10 | 28.309747 |
| **COC17** | 12451928 | 12011070 | 96.81 | 0.0851895 | 0.940585457 | 0.825846603 | 7 | 23.215236 |
| **COC19** | 15125859 | 14643166 | 97.19 | 0.0867262 | 0.941148308 | 0.874656605 | 9 | 26.718768 |
| **COC7** | 8533452 | 8025088 | 84.32 | 0.17473 | 0.883801574 | 0.5352049 | 4 | 14.979581 |
| **HOR16** | 20026317 | 19410958 | 97.33 | 0.0795994 | 0.945680497 | 0.914645713 | 14 | 39.108156 |
| **HOR4** | 17530713 | 17103746 | 97.89 | 0.0925175 | 0.942631583 | 0.881709173 | 10 | 28.726877 |
| **HOR6** | 19567040 | 18954168 | 97.25 | 0.0799675 | 0.946704167 | 0.918558619 | 12 | 34.671127 |
| **HOR7** | 16536756 | 16049354 | 97.36 | 0.101436 | 0.939901849 | 0.855900735 | 8 | 25.653929 |
| **MUR15** | 19893668 | 19235198 | 97 | 0.0966443 | 0.935523187 | 0.857908363 | 10 | 29.936208 |
| **MUR16** | 12603765 | 12159944 | 96.88 | 0.0774015 | 0.953907444 | 0.824251475 | 7 | 24.973462 |
| **MUR17** | 15669733 | 14511434 | 92.83 | 0.122035 | 0.922430254 | 0.776049139 | 7 | 21.85571 |
| **MUR20** | 14235087 | 13861624 | 97.75 | 0.100851 | 0.935910925 | 0.859243787 | 9 | 26.8259 |
| **PAL12** | 18863248 | 19341898 | 97.55 | 0.082103 | 0.946568051 | 0.900930129 | 10 | 28.776939 |
| **PAL16** | 3958526 | 3810546 | 96.63 | 0.213877 | 0.837400308 | 0.254548432 | 2 | 9.055073 |
| **PAL6** | 8662644 | 8307962 | 96.28 | 0.110427 | 0.921523735 | 0.673732957 | 5 | 17.715713 |
| **PAL7** | 12545857 | 12126038 | 97.03 | 0.0853889 | 0.943596279 | 0.846446533 | 7 | 23.099512 |
| **PAN1** | 13493783 | 13074414 | 97.22 | 0.0961753 | 0.93322509 | 0.788037511 | 7 | 22.18564 |

| Line | Total Reads | Mapped Reads paired | Mapping Rate paired | Frequency Missing Sites | Percent 1X | Percent 4X | Median Depth | Average Quality (NA excluded) |
|---|---|---|---|---|---|---|---|---|
| **PAN5** | 11454217 | 10947794 | 95.78 | 0.171477 | 0.89062912 | 0.557564965 | 4 | 14.332644 |
| **PAN9** | 14462860 | 13966798 | 96.9 | 0.0943379 | 0.939613869 | 0.853602381 | 8 | 25.011322 |
| **PIN3** | 17256729 | 16786270 | 97.61 | 0.0873108 | 0.94239514 | 0.886942336 | 10 | 28.245879 |
| **PIN6** | 12279097 | 11842198 | 96.83 | 0.0839895 | 0.943469274 | 0.863768506 | 8 | 24.737327 |
| **PIN7** | 14845446 | 14423300 | 97.5 | 0.0913857 | 0.939771193 | 0.855498068 | 8 | 24.938503 |
| **PIN9** | 19513663 | 18921394 | 97.28 | 0.0785065 | 0.944568986 | 0.891827492 | 10 | 30.584467 |
| **POB10** | 17333118 | 16897498 | 97.77 | 0.113248 | 0.93374925 | 0.839942669 | 8 | 24.433195 |
| **POB16** | 15815482 | 15285062 | 97.06 | 0.084191 | 0.945719821 | 0.899785275 | 10 | 30.130401 |
| **POB19** | 29069830 | 28034352 | 96.81 | 0.0788636 | 0.949081053 | 0.931900094 | 17 | 45.177738 |
| **POB7** | 12272121 | 11860176 | 97.01 | 0.0914374 | 0.938619091 | 0.803773907 | 7 | 22.173469 |
| **RAB17** | 20142894 | 19644904 | 97.89 | 0.0867714 | 0.942018708 | 0.912441866 | 12 | 34.761648 |
| **RAB20** | 25499341 | 24844970 | 97.76 | 0.083803 | 0.943951032 | 0.920764855 | 15 | 40.132486 |
| **RAB4** | 5574554 | 5372236 | 96.67 | 0.246698 | 0.822850644 | 0.373876352 | 3 | 11.745683 |
| **RAB9** | 17813030 | 17252450 | 97.26 | 0.076267 | 0.949048009 | 0.921511878 | 12 | 35.008764 |
| **SPE2** | 14723146 | 14292882 | 97.44 | 0.0907326 | 0.93888785 | 0.856997056 | 8 | 26.091352 |
| **SPE5** | 9009297 | 8542706 | 95.02 | 0.259712 | 0.836739117 | 0.482691684 | 3 | 13.762534 |
| **SPE6** | 21149166 | 20476064 | 97.19 | 0.0774476 | 0.948390664 | 0.912880539 | 13 | 36.429486 |
| **SPE7** | 10718970 | 10455162 | 97.92 | 0.10757 | 0.930315283 | 0.777761151 | 6 | 20.87309 |
| **VDM17** | 17074008 | 16566130 | 97.37 | 0.0873811 | 0.940959966 | 0.883627653 | 9 | 28.295409 |
| **VDM20** | 18009082 | 16097200 | 89.71 | 0.0839738 | 0.944618675 | 0.88943338 | 10 | 29.29148 |
| **VDM9** | 11975809 | 11616758 | 97.33 | 0.110689 | 0.920913962 | 0.728168666 | 6 | 20.901281 |
| **VIE16** | 14678603 | 14160160 | 96.79 | 0.0841149 | 0.941838295 | 0.846866663 | 8 | 24.412579 |
| **VIE3** | 24454713 | 23797668 | 97.58 | 0.0858097 | 0.943619342 | 0.890739707 | 11 | 31.334127 |
| **VIE4** | 15045325 | 14539938 | 96.95 | 0.091256 | 0.941023572 | 0.844092725 | 8 | 23.671476 |
| **VIE6** | 17529193 | 17034084 | 97.54 | 0.0823222 | 0.944623981 | 0.906702361 | 11 | 31.234305 |

| Line | Total Reads | Mapped Reads paired | Mapping Rate paired | Frequency Missing Sites | Percent 1X | Percent 4X | Median Depth | Average Quality (NA excluded) |
|---|---|---|---|---|---|---|---|---|
| **mean** | 15753452 | 15224790 | 96.58 | 0.113558 | 0.924525049 | 0.792506667 | 8.451 | 25.645610 |
| **median** | 15397796 | 14591552 | 97.115 | 0.088198 | 0.940772712 | 0.856448895 | 8 | 25.232966 |
| **sum** | 976714024 | 943936960 | | | | | | |

**Table S3. Model Summary Table**. All models include the centromere. All flowers phenotyped (Full) or only flowers from the sequenced genotypes (Seq) denoted. Adding a quadratic elevation term increased adjusted $r^2$ when predicting population mean short stamen number, but a quadratic elevation term did not increase model accuracy when predicting nucleotide diversity and a quadratic nucleotide diversity term did not increase model accuracy compared to only a linear nucleotide diversity term. Residuals of model 4 are used as response variable in model 8. Residuals of model 6 are used as the response variable in model 7.

| | **Response Variable** | Elevation estimate | Elevation p value | centered Elevation$^2$ estimate | centered Elevation$^2$ p value | Nucleotide Diversity estimate | Nucleotide Diversity p value | $r^2$ | adjusted $r^2$ | AIC score |
|---|---|---|---|---|---|---|---|---|---|---|
| 1 | Nucleotide Diversity | -1.81E-06 | **0.00019** | | | | | 0.6411 | 0.6155 | -180.67 |
| 2 | Full_PopFlwrMean | 3.86E-04 | **0.0107** | | | | | 0.3819 | 0.3377 | 8.01379 |
| 3 | Full_PopFlwrMean | 4.82E-04 | **0.0041** | -4.73E-07 | 0.137267 | | | 0.4818 | 0.4021 | 7.19102 |
| 4 | Seq_PopFlwrMean | | | | | -61.5985 | 0.387 | 0.05398 | -0.0359 | 13.4079 |
| 5 | Seq_PopFlwrMean | 6.47E-04 | **0.00882** | -2.99E-07 | 0.34514 | 1.39E+02 | 0.16696 | 0.4886 | 0.3608 | 7.56529 |
| 6 | Seq_PopFlwrMean | 4.18E-04 | **0.01** | -4.07E-07 | 0.2101 | | | 0.3964 | 0.3035 | 8.21867 |
| 7 | Short Stamen Residuals | | | | | 4.68E+01 | 0.397 | 0.05168 | -0.0161 | 5.36963 |
| 8 | Short Stamen Residuals | 3.17E-04 | 0.0613 | -4.54E-07 | 0.1955 | | | 0.2559 | 0.1414 | 10.6789 |

**Table S4. Shared hits from short stamen number GWAS**. All GWAS conducted with GEMMA. P-values are correlated with effect sizes in all GWAS. Chr, Window, and Position indicate the chromosomal position of each SNP. For each GWAS, the effect size (beta), standard error of the effect size (SE), and p-value are reported. The effect sizes represent the difference in mean short stamen number from one allele being replaced. For the arcsine transformed GWAS, effect sizes and standard errors have not been back transformed. In the binary GWAS, the sign of the effect size is reversed because no short stamen loss (i.e., mean short stamen number = 2) was coded as a zero and any short stamen loss (i.e., mean short stamen number < 2) was coded as a one. The second page repeats the first 3 columns and reports if the SNP is in a gene, the chromosomal position of the gene, and the gene type from GBrowse with TAIR10 annotation. Notes information is from annotations within GBrowse or literature search of papers that reference each gene as reported by GBrowse, cited within the table.

| Shared Short Stamen | | | Arcsine Transformed | | | Raw Phenotype | | | Binary: any loss? | | | Subset: if some loss | | |
|---|---|---|---|---|---|---|---|---|---|---|---|---|---|---|
| **Chr** | **Window** | **Position** | **Beta** | **SE** | **p** | **Beta** | **SE** | **p** | **Beta** | **SE** | **p** | **Beta** | **SE** | **p** |
| 1 | A | 14262517 | | | | -0.33 | 0.04 | 1.18E-11 | | | | -0.30 | 0.04 | 1.25E-09 |
| 3 | A | 2239234 | 0.21 | 0.04 | 3.08E-07 | | | | -0.28 | 0.06 | 2.06E-05 | | | |
| | B | 2253161 | -0.22 | 0.04 | 1.19E-07 | | | | 0.28 | 0.06 | 1.27E-05 | | | |
| | C | 2942726 | -0.22 | 0.03 | 1.21E-08 | -0.26 | 0.05 | 4.42E-07 | 0.25 | 0.05 | 1.64E-05 | | | |
| 5 | A | 4899729 | | | | | | | 0.23 | 0.06 | 1.64E-04 | | | |
| | | 4899733 | | | | | | | 0.23 | 0.06 | 1.64E-04 | | | |
| | | 4899789 | | | | | | | 0.24 | 0.06 | 9.33E-05 | | | |
| | | 4899798 | -0.26 | 0.05 | 4.66E-07 | -0.33 | 0.06 | 9.49E-07 | 0.27 | 0.06 | 2.98E-05 | | | |
| | | 4899803 | -0.26 | 0.05 | 4.66E-07 | -0.33 | 0.06 | 9.49E-07 | 0.27 | 0.06 | 2.98E-05 | | | |
| | | 4899865 | | | | | | | 0.23 | 0.06 | 1.64E-04 | | | |
| | | 4900275 | | | | | | | 0.25 | 0.06 | 5.11E-05 | | | |
| | | 4900560 | | | | | | | 0.24 | 0.05 | 2.58E-05 | | | |
| | | 4900628 | | | | | | | 0.23 | 0.06 | 1.72E-04 | | | |
| | B | 4919727 | | | | | | | 0.24 | 0.06 | 8.00E-05 | | | |
| | B,C | 4920179 | -0.24 | 0.04 | 1.02E-07 | | | | 0.32 | 0.06 | 4.96E-06 | | | |
| | C | 4920289 | | | | | | | 0.24 | 0.05 | 4.49E-05 | | | |
| | | 4920304 | | | | | | | 0.24 | 0.05 | 4.49E-05 | | | |
| | | 4920731 | | | | | | | 0.25 | 0.05 | 1.87E-05 | | | |
| | D | 10731997 | 0.20 | 0.03 | 3.11E-07 | 0.23 | 0.04 | 2.47E-06 | -0.28 | 0.07 | 9.62E-05 | | | |
| | E | 13458838 | -0.19 | 0.03 | 4.48E-08 | -0.24 | 0.03 | 4.59E-09 | | | | -0.26 | 0.04 | 4.98E-07 |

| Shared Short Stamen | | | | | | |
|---|---|---|---|---|---|---|
| **Chr** | **Window** | **Position** | **In a gene?** | **Gene Position** | **Gene Type** | **notes** |
| 1 | A | 14262517 | AT1TE46795 | Chr1:14262180..14263528 (+ strand) | transposable_element | in the centromere |
| 3 | A | 2239234 | AT3G07070.1 | Chr3:2237897..2240266 (+ strand) | Protein kinase superfamily protein (PBL26, PBS1-LIKE 26) | near a GTP-binding family protein required for maintenance of inflorescence meristem identity, floral organ development, and megaspore mother cell specification |
| | B | 2253161 | AT3G07110 | Chr3:2251946..2253597 (+ strand) | Ribosomal protein L13 family protein | expressed in stamen (and all other plant parts). About 40kb away from a SMAD/FHA domain-containing protein (Forkhead domain protein that is a subunit of ISWI chromatin remodeling complex.Interacts with histones and regulates the expression of genes involved in stamen filament elongation.) |
| | C | 2942726 | AT3G09580.1 | Chr3:2942158..2944457 (- strand) | FAD/NAD(P)-binding oxidoreductase family protein | Expressed in stamen (and all other plant parts). Oxidoreductase located in chloroplast. |
| 5 | A | 4899729 | DUF1637; AT5G15120 | Chr5:4898641..4900734 (+ strand) | HUP29, hypoxia response unknown protein 29, PCO1, Plant cysteine oxidase 2 | involved in anaerobic respiration, cellular response to hypoxia, and peptidyl-cysteine oxidation; (Mustroph et al., 2010 Plant Physiology); Is near AT5G15110 which is expressed in pollen |
| | | 4899733 | | | | |
| | | 4899789 | | | | |
| | | 4899798 | | | | |
| | | 4899803 | | | | |
| | | 4899865 | | | | |
| | | 4900275 | | | | |
| | | 4900560 | | | | |
| | | 4900628 | | | | |
| | B | 4919727 | No | Chr5:4919548..4919824 (+ strand) | long non-coding RNA | |
| | B,C | 4920179 | No | Chr5:4919989..4919989 | TE | |
| | C | 4920289 | No | Chr5:4920043..4920718 (+ strand) | long non-coding RNA | |
| | | 4920304 | AT5TE17795 | Chr5:4920665..4921403 (+ | TE | |
| | | 4920731 | | | | |
| | D | 10731997 | AT5G28692 | Chr5:10728378..10733393 (- strand) | transposable_element_gene; locus:504955064 | gypsy-like retrotransposon family |
| | E | 13458838 | No | | | closest thing: AT5G35200 a ENTH/ANTH/VHS superfamiliy protein (PICALM3) - PICALM 5 and 5b are associated with pollen tube growth (Muro et al. 2018 Commun Biol.) |
